# Stabilizing role of seed banks and the maintenance of bacterial diversity

**DOI:** 10.1101/2020.10.05.327387

**Authors:** Nathan I. Wisnoski, Jay T. Lennon

## Abstract

Coexisting species often exhibit negative frequency dependence due to mechanisms that promote population growth and persistence when rare. These stabilizing mechanisms can maintain diversity through interspecific niche differences, but also through life-history strategies like dormancy that buffer populations in fluctuating environments. However, there are few tests demonstrating how seed banks contribute to long-term community dynamics and the maintenance of diversity. Using a multi-year, high-frequency time series of bacterial community data from a north temperate lake, we documented patterns consistent with stabilizing coexistence. Bacterial taxa exhibited differential responses to seasonal environmental conditions, while seed bank dynamics helped maintain diversity over winter. Strong negative frequency dependence in rare, but metabolically active, taxa suggested a role for biotic interactions in promoting coexistence. Together, our results provide field-based evidence that niche differences and seed banks contribute to recurring community dynamics and the long-term maintenance of diversity in nature.

## INTRODUCTION

The maintenance of biodiversity is important for regulating species interactions, stabilizing ecosystem functions, and promoting resilience in response to perturbations. Diversity is maintained by many processes, including niche differentiation in resource use (Tilman 1982; Gudelj *et al.* 2010; Johnson *et al.* 2012), defensive abilities (Leibold 1996; Thingstad *et al.* 2014; Cadier *et al.* 2019), and abiotic constraints (Holt 2009). These stabilizing niche differences among species contribute to the maintenance of diversity by causing species to limit their own growth more than the growth of other species, thereby preventing competitive exclusion and allowing populations to recover from low abundances (Chesson 2000; Chase & Leibold 2003; Adler *et al.* 2007). Some stabilizing mechanisms of coexistence rely on environmental fluctuations and can further increase the number of species in a community (Chesson 1994; Chesson & Huntly 1997; Descamps-Julien & Gonzalez 2005). For example, in seasonal environments, species that are favored at different times of the year may be able to coexist in the community if they can survive through periods of unfavorable environmental conditions (Pake & Venable 1996). Given the near ubiquity of environmental variability, a central and unresolved question is how stabilizing mechanisms promote the maintenance of diversity across the wide range of taxonomic groups and ecosystems that exist in nature.

Stabilization from niche differences should generate negative frequency-dependence (NFD) in population growth. The implication of NFD is that rare populations grow faster than common populations (Chesson 2000; Adler *et al.* 2007). NFD may arise from mechanisms that promote coexistence in relatively constant environments, such as trade-offs in resource acquisition and allocation, but also in temporally variable environments. In fluctuating environments, the storage effect is a coexistence mechanism that reflects the ability of species to grow well during favorable conditions while minimizing losses during unfavorable conditions by “storing” individuals in long-lived life stages (Warner & Chesson 1985; Chesson 2000). The storage effect requires that taxa differ in their responses to environmental conditions, that intraspecific limitation peaks during favorable conditions, and that population growth is buffered in suboptimal environments (Pake & Venable 1996; Cáceres 1997; Angert *et al.* 2009). The storage effect may be particularly important in communities where species experience periods of extremely slow growth (Gray *et al.* 2019) or engage in various forms of dormancy, which are common among plants and animals, but also microorganisms (Lennon & Jones 2011).

Support for stabilizing coexistence has largely come from plant and animal communities (Cáceres 1997; Angert *et al.* 2009; Yenni *et al.* 2017), while evidence for its role in complex microbial systems is less common (Zhang *et al.* 2010). In microbial communities, which contain a disproportionately large number of rare taxa (Sogin *et al.* 2006; Lynch & Neufeld 2015; Shade *et al.* 2018), populations may vary widely in their stability and long-term contributions to diversity. Although rare taxa are prone to extinction (Lande 1993), some persist for longer periods of time (Alonso-Sáez *et al.* 2015; Lynch & Neufeld 2015; Newton & Shade 2016). In fluctuating environments, many of these rare taxa can quickly respond to favorable conditions (Shade *et al.* 2014; Linz *et al.* 2017; Nyirabuhoro *et al.* 2020), suggesting that temporally variable opportunities for growth may be important for population persistence. While niche differences among bacterial taxa are well documented (Lennon *et al.* 2012; Evans *et al.* 2014; Meier *et al.* 2017), the long-term implications of these differences and their contribution to the maintenance of biodiversity in nature remain understudied.

A leading hypothesis for the long-term maintenance of microbial diversity has been coexistence mediated by dormant seed banks (Jones & Lennon 2010; Lennon & Jones 2011; Mestre & Höfer 2020; Sorensen & Shade 2020). Dormancy can buffer microbial populations in different ways (Rittershaus *et al.* 2013). For example, some species form physical resting structures that protect individuals from harsh abiotic stress (Setlow 2006; de Rezende *et al.* 2013). Other species may reduce mortality associated with resource limitations by shifting energetic demands from growth to maintenance energy levels (Lennon & Jones 2011; Hoehler & Jørgensen 2013; Lever *et al.* 2015). Dormancy may even protect against top-down pressure from grazers (which may be unable to digest or extract energy from starved cells or endospores) and phage (which cannot replicate due to inactive host machinery) (Pernthaler 2005; Klobutcher *et al.* 2006; Bautista *et al.* 2015; Kearney *et al.* 2018). Consequently, dormant bacteria may exhibit reduced mortality in the environment (Hoehler & Jørgensen 2013), thereby accumulating into seed banks until favorable conditions return (Wörmer *et al.* 2019). Much insight has been gained from short-term microbial studies and analogies with plant and zooplankton communities, but evidence from long-term field studied demonstrating how the temporal dynamics of microbial seed banks help maintain diversity in fluctuating environments is lacking.

In this study, we tracked bacterioplankton dynamics over time in a north temperate lake using high-resolution molecular data to infer ecological processes that maintain microbial diversity. Bacterial communities in fluctuating aquatic environments often exhibit recurrent, seasonal community patterns (Shade *et al.* 2007; Gilbert *et al.* 2012; Fuhrman *et al.* 2015; Ward *et al.* 2017), but the potential mechanisms that contribute to cyclical dynamics and maintain diversity in nature are poorly resolved. We characterized how persistent (and putatively coexisting) taxa respond to environmental fluctuations and used null models to assess whether stabilizing biotic interactions (e.g., self-limitation, as evidenced by strong NFD) help maintain rare, but metabolically active, taxa in the community (Yenni *et al.* 2017; Rovere & Fox 2019).

Specifically, we compared patterns of NFD and population dynamics in the active and total portions of the community (inferred by 16S rRNA transcripts and genes, respectively) to quantify the importance of slow growth or dormancy strategies for the maintenance of diversity. Our results provide empirical evidence that stabilizing biotic interactions and seed bank dynamics underlie seasonal community dynamics and play key roles in maintaining bacterial diversity in natural ecosystems.

## METHODS

### Study site and sampling

University Lake is a 3.2 ha meso-eutrophic reservoir located in the Indiana University Research and Teaching Preserve, Bloomington, Indiana, USA (39°11’ N, 86°30’ W). The surrounding watershed is dominated by oak, beech, and maple forests. Three streams drain into University Lake, which has an estimated volume of 150,000 m^3^ and a maximum depth of 10 m. From April 2013 to September 2015, we took weekly water samples (1 L) from the epilimnion using a 1 m depth-integrated sampler for microbial biomass, and measured environmental variables commonly associated with aquatic microbial community dynamics: total phosphorus (TP), total nitrogen (TN), and dissolved organic carbon (DOC). Microbial biomass was filtered on 0.2 μm Supor filters (Pall, Port Washington, NY, USA) and frozen at −80 °C. We quantified TP using the ammonium molybdate method (Wetzel & Likens 2000) and TN with the second derivative method after persulfate digestion (Crumpton *et al.* 1992). DOC was quantified on 0.7 μm filtrates using nondispersive infrared (NDIR) detection on a Shimadzu TOC-V (Kyoto, Japan). We also quantified water transparency with a Secchi disk and used a Quanta Hydrolab (OTT, Kempton, Germany) water sonde to measure temperature, conductivity, dissolved oxygen, salinity, and pH of the samples.

### Bacterial community structure

We characterized the structure of the bacterial community using high-throughput 16S rRNA sequencing. We extracted total nucleic acids from biomass retained on 0.2 μm filters using the MoBio PowerWater RNA extraction kit and the DNA elution accessory kit. Because sequences obtained from DNA can come from metabolically active or inactive (e.g., slow growing or dormant) individuals, this sample represents the “total” community. In contrast, RNA is a more ephemeral molecule that is essential for synthesizing proteins; therefore, it is often used to characterize the metabolically “active” subset of the community (Molin & Givskov 1999; Steiner *et al.* 2019; Locey *et al.* 2020). After extracting nucleic acids, we used DNase (Invitrogen) to remove DNA from the RNA extractions and then synthesized cDNA with SuperScript III First Strand Synthesis kit and random hexamer primers (Invitrogen). To amplify the 16S rRNA gene (DNA) and transcripts (cDNA), we used barcoded V4 primers (515F and 806R) designed for the Illumina MiSeq platform (Caporaso *et al.* 2012). We then purified the PCR products with AMPure XP, quantified DNA concentrations using PicoGreen, and pooled samples at 10 ng per sample. The resulting libraries were sequenced on an Illumina MiSeq at the Indiana University Center for Genomic and Bioinformatics Sequencing Facility using 250 × 250 bp paired-end reads (Reagent Kit v2). Sequences were subsequently processed using the software package mothur (version 1.41.1) (Schloss *et al.* 2009). We assembled contigs, removed low quality sequences (minimum score of 35), aligned sequences to the SILVA Database (version 132) (Quast *et al.* 2013), removed chimeras using the VSEARCH algorithm (Rognes *et al.* 2016), and created 97% similar operational taxonomic units (OTUs) using the OptiClust algorithm (Westcott & Schloss 2017), and classified sequences with the RDP taxonomy (Cole *et al.* 2009). To account for variation in sequencing depth, subsequent analyses were performed on rarefied abundance data subsampled to the fewest number of reads in the time series (*N* = 5979 per sample) using R (version 3.6.0) (R Core Team 2020).

### Differential responses to environment

We evaluated whether niche partitioning occurred along a suite of environmental variables. First, we performed a principal component analysis (PCA) on Hellinger-transformed abundances to visualize seasonal patterns of compositional trajectories. Next, we identified environmental drivers of community dynamics using redundancy analysis (RDA). We then looked more closely at the persistent, and potentially coexisting, subset of taxa in the community (OTUs present in ≥80% of the DNA-based samples, Table S1). To determine whether environmental fluctuations facilitated temporal niche partitioning, we identified the week of the year when each persistent OTU (n = 82) experienced its maximum average growth rate (see calculations in *Stabilizing niche differences* below). For each of the persistent OTUs, we compared the environmental conditions at the time of the year when they experienced maximum growth (as an index of optimal conditions along an annual environmental cycle). Specifically, we performed a principal component analysis (PCA) on the environmental variables including temperature, specific conductivity, transparency, pH, TP, TN, and DOC (all standardized to mean = 0, standard deviation = 1). We plotted each time point along the first two PC axes, along with the PC loadings of the environmental variables, and labeled each point with the name of the OTU that exhibited maximum growth at that particular time point (Fig. S1).

### Stabilizing niche differences

We then inferred whether niche differences contributed to patterns of negative frequency dependence (NFD) in the community. In particular, we examined (1) whether growth exhibited negative frequency dependence overall, which is indicative of stabilization in the community, and (2) whether rare taxa experienced stronger negative frequency dependence than common taxa (Yenni *et al.* 2012, 2017; Rovere & Fox 2019). Such patterns of NFD shed light on coexistence for three reasons. First, taxa are more likely to be common in the community when they are environmentally favored relative to other taxa in the community. Second, taxa that experience strong intraspecific limitation during environmentally favorable periods should grow faster when rare than when common, thereby generating NFD. Third, differences among taxa in the strength of self-limitation during favored growth periods should lead to different average relative abundances in the community, implying that rare yet persistent taxa may be strongly stabilized (Yenni *et al.* 2012, 2017; Rovere & Fox 2019). Patterns of NFD often arise from density-dependent processes, such as nutrient limitation, but density dependence will only generate NFD if species limit their own growth more strongly than they limit the growth of other species (Adler *et al.* 2007). However, metabolically inactive individuals are unlikely to engage in the biotic interactions that generate NFD. Therefore, we focused on NFD in the metabolically active (i.e., RNA-based) portion of the community comprised of the persistent taxa (Table S1).

We then calculated NFD for each OTU by comparing rates of change in relative abundance between weekly samples. We inferred the strength of NFD for a given OTU as the magnitude of the negative slope of the relationship between an OTU’s relative abundance and its per capita growth rate at each time step (*t*) across the time series. We calculated the relative abundance (*x*_*t,s*_), of each OTU (*s*) as its abundance (*N*_*t,s*_) in the community of *s* OTUs relative to the total abundance of all *s* OTUs (∑_*s*_ *N*_*t,s*_) at a given time step (*t*), such that 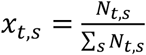. From this, we then calculated the natural log of the per capita growth rate of each OTU as 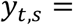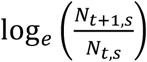. To estimate the strength of NFD for each OTU, we fit simple linear regressions (*y*_*s*_ = *β*_0,*s*_ + *β*_1,*s*_*x*_*s*_ + *∈*_*s*_), where the equilibrium frequency of an OTU (*f*) is the x-intercept, 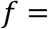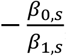, and the degree of NFD is the slope, NFD = *β*_1,*s*_. In the end, *f* describes whether an OTU is common or rare, and negative slopes with greater magnitudes indicate stronger negative frequency dependence.

### Stabilization of rare bacterial taxa

We then inferred whether rare taxa exhibited stronger NFD than expected by chance. Stabilization of rare taxa would be supported if OTUs with lower equilibrium frequencies (smaller values of *f*) had more negative slopes (larger |*β*_1,*s*_|) indicating stronger self-limitation. We inferred the overall relationship between rarity and strength of NFD from the covariance between the log(NFD) and log(*f*): a more negative covariance would indicate that rarer taxa were more strongly stabilized than common taxa. To account for the fact that the expectation of this covariance is already negative, and to control for spurious statistical correlations in the temporal data due to other factors, we implemented a null model approach (Yenni *et al.* 2017; Rovere & Fox 2019). We shuffled the abundances of each OTU independently, recalculated relative abundances and per capita growth rates, estimated equilibrium frequencies (*f*s) and negative frequency dependences (NFDs), and calculated the covariance, repeating this procedure 5000 times to generate a null distribution of covariance values (*COV*[log(*f*), log(NFD)]) (Yenni *et al.* 2017). This procedure maintains the potential abundances detected for each OTU but erases the temporal structure of each taxon’s growth dynamics, removing signatures of intraspecific limitation as well as any interspecific limitations correlated with the population dynamics of other OTUs. We then compared our observed covariance with the null distribution to infer the strength of asymmetry in NFD (i.e., the degree to which rare OTUs experience disproportionately stronger self-limitation than common OTUs). We quantified divergence from null distributions using standardized effect sizes (SES = mean observed covariance / standard deviation of covariances in the null distribution) and the ratio of observed covariance to the average covariance of the null distribution (Yenni *et al.* 2017). More negative SES values and larger ratios would indicate greater deviations from the null expectation of equal NFD across taxa. We inferred the degree of statistical significance by calculating a p-value as the proportion of null covariance values less than or equal to our observed covariance.

### Seed bank dynamics

Given the hypothesis that seed banks are important for the maintenance of bacterial diversity in nature, we analyzed the temporal dynamics of buffered population growth, a key criterion of the storage effect. First, we examined whether the seed bank served as a reservoir of taxonomic diversity by comparing the ratio of total richness to active richness at each time point in the time series, where larger ratios indicate that the total community had higher α-diversity than the active subset of the community. Second, we sought to determine whether seed bank dynamics were more important for the maintenance of rare or common taxa in the community. To do so, we developed a reactivation metric to quantify each OTU’s frequency of reactivation from the seed bank. For each OTU, its reactivation score is the number of times an OTU was present (i.e., detected in the DNA pool) but likely in an inactive (i.e., absent from the RNA pool) state at time point *t*, yet active (present in the RNA pool) at the subsequent time point *t+1*. This represents a transition from the inactive to active state mediated by slow growth or dormancy based on recovery of sequences in the DNA and RNA pools. Thus, OTUs with higher reactivation scores may be more reliant on the seed bank for long-term persistence in the community. We then analyzed the relationship between the average relative abundance of active OTUs (excluding zeroes) and their reactivation score to determine whether seed banking was more important for maintaining rare taxa than common taxa in the bacterioplankton community.

We compared observed patterns of reactivation to null models of community dynamics because the probability of resuscitation may not be independent of relative abundance. Such non-independence could be expected, for example, if rarer OTUs were more likely to be inactive than common OTUs. We generated null models of bacterial time series (n = 1000) by randomly redistributing observed counts of each OTU across the time series, keeping total observed counts for each OTU constant to preserve the relationships among common and rare taxa in the community. By redistributing individuals across sampling dates for each taxon independently, we removed population dynamic signatures of intraspecific density dependence as well as interspecific density dependence arising from biotic interactions with other taxa in the community. Thus, our null models represent the range of reactivation scores possible for an OTU of a given mean relative abundance in the community if its dynamics were stochastic. We then compared our observed reactivation scores to the null models to identify whether common or rare OTUs reactivated more or less frequently than expected by chance.

## RESULTS

### Differential responses to environment

Bacterial community dynamics were related to environmental variability, with different taxa favored at different times of the year. During the summer months, the community followed a recurrent successional trajectory (Fig. 1A). This trajectory was strongly aligned with seasonal trends in temperature (Fig. 1B). Across longer time scales, inter-annual variation in dissolved oxygen and pH was associated with compositional differences in the active bacterial community during winter months. Within an annual cycle, the persistent OTUs (n = 82) demonstrated temporal partitioning in their maximal growth rates in the active portion of the community (Fig. 2, Table S1), corresponding to different environmental conditions (Fig. S1).

**Figure 1.**
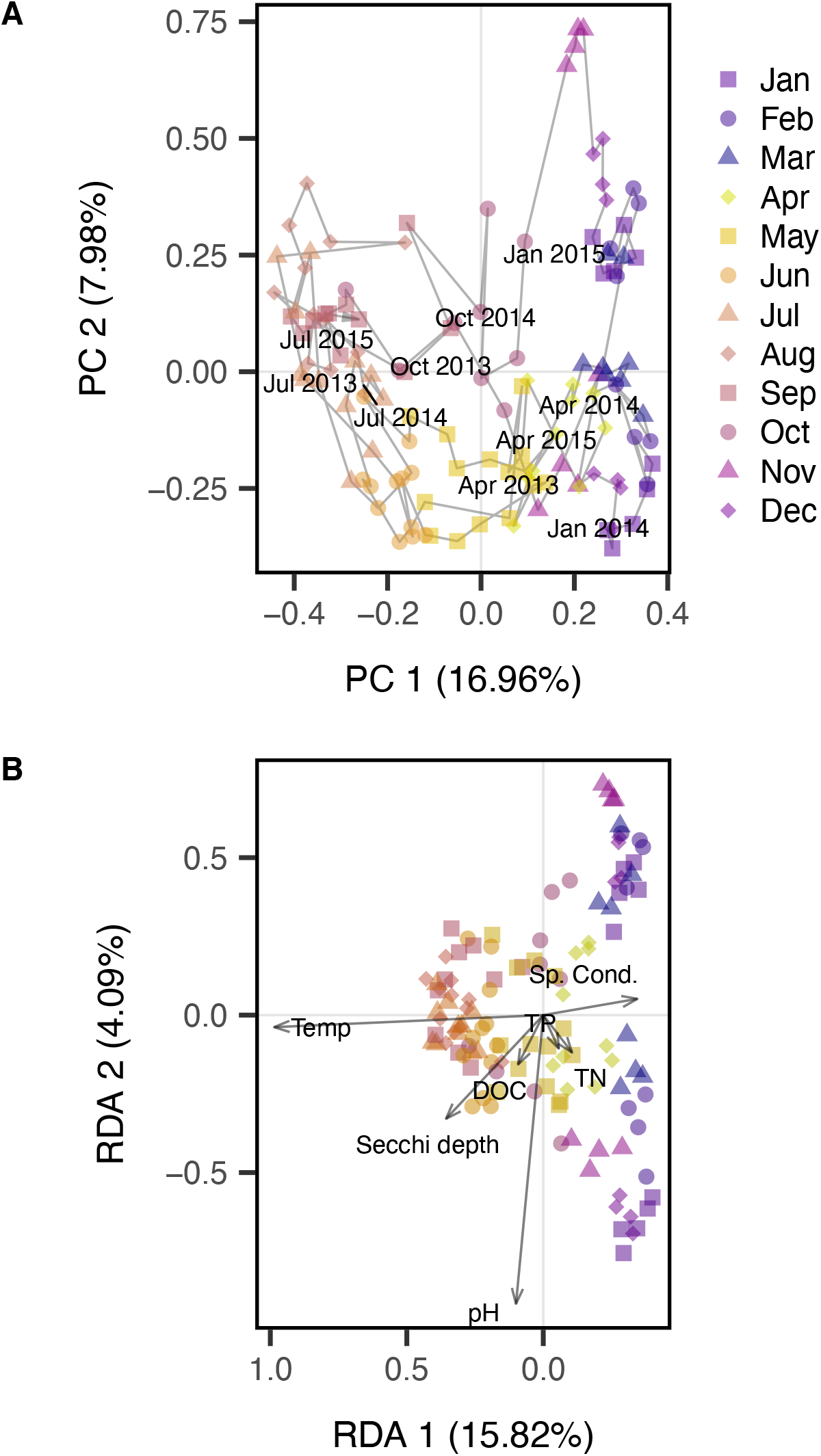
Seasonal dynamics of the active bacterial community in University Lake. (A) The compositional trajectory of the active community (determined by high-throughput sequencing of 16S rRNA transcripts) shows strong seasonality, but the community remains relatively static over winter. The first two axes of the principal component analysis (PCA) depict summer/winter differences (PC1) along the major axis, and slight inter-annual differences in winter composition (cool colors) along the minor axis (PC2). The summer successional trajectory (warmer colors) is highly repeatable across years. (B) Constrained ordination using redundancy analysis (RDA) shows the environmental drivers of community structure, along with strong correlates of individual taxa in the community. This analysis reveals that differences in pH explain variation in winter composition among years.

**Figure 2.**
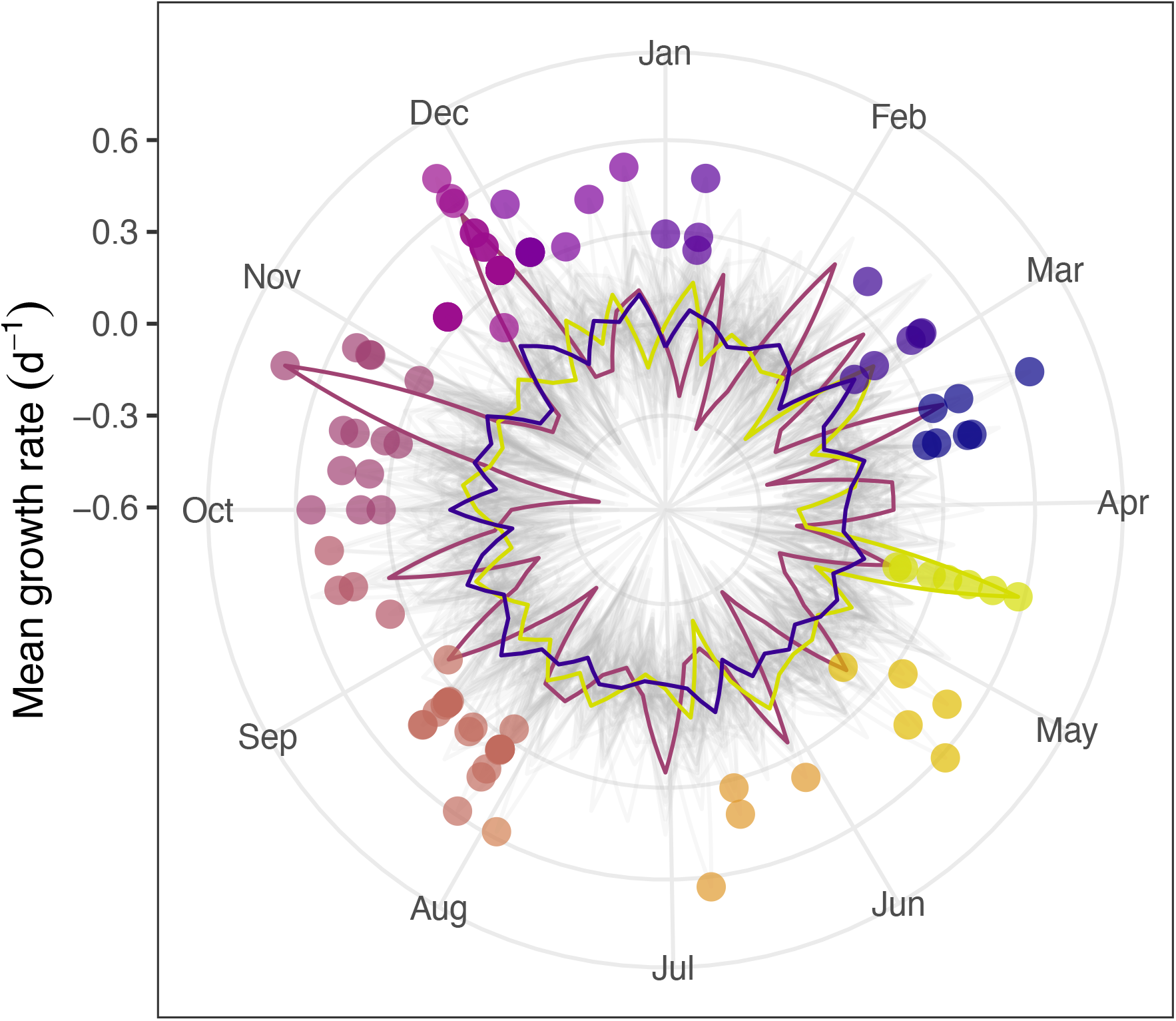
Temporal partitioning of maximum growth rate among persistent bacterial taxa in University Lake. Lines represent the mean daily growth rate for each taxon over the time series. Points indicate the maximum growth observed for each bacterial taxon (OTU). Overall, the 82 persistent OTUs have maximum growth rates at different seasons of the year. Points are color-coded such that warmer colors correspond to spring and summer months and cooler colors correspond to winter months. Colored lined trace out the growth dynamics of three individual taxa with different environmental responses (blue = OTU 1, Betaproteobacteria; yellow = OTU 17, Actinobacteria; mauve = OTU 18, Gammaproteobacteria). More taxonomic details can be found in Table S1.

### Stabilizing biotic interactions

Persistent taxa exhibited stabilizing NFD, which varied in strength depending on each taxon’s mean relative abundance in the community (Fig. S2). In particular, NFD was significantly stronger for rare taxa than common taxa, but only in the active portion of the community (p = 0.0002; SES = −4.03, covariance ratio = 1.08), not in the total community (p = 0.221; SES = −0.777, covariance ratio = 1.01) (Fig. 3). The p-values reflect the rank of observed NFD compared with null simulations, while SES values take into account the variance in the null distribution. In other words, the total community showed nearly the same degree of stabilization (|SES| < 2, covariance ratio ~ 1) as the null communities.

**Figure 3.**
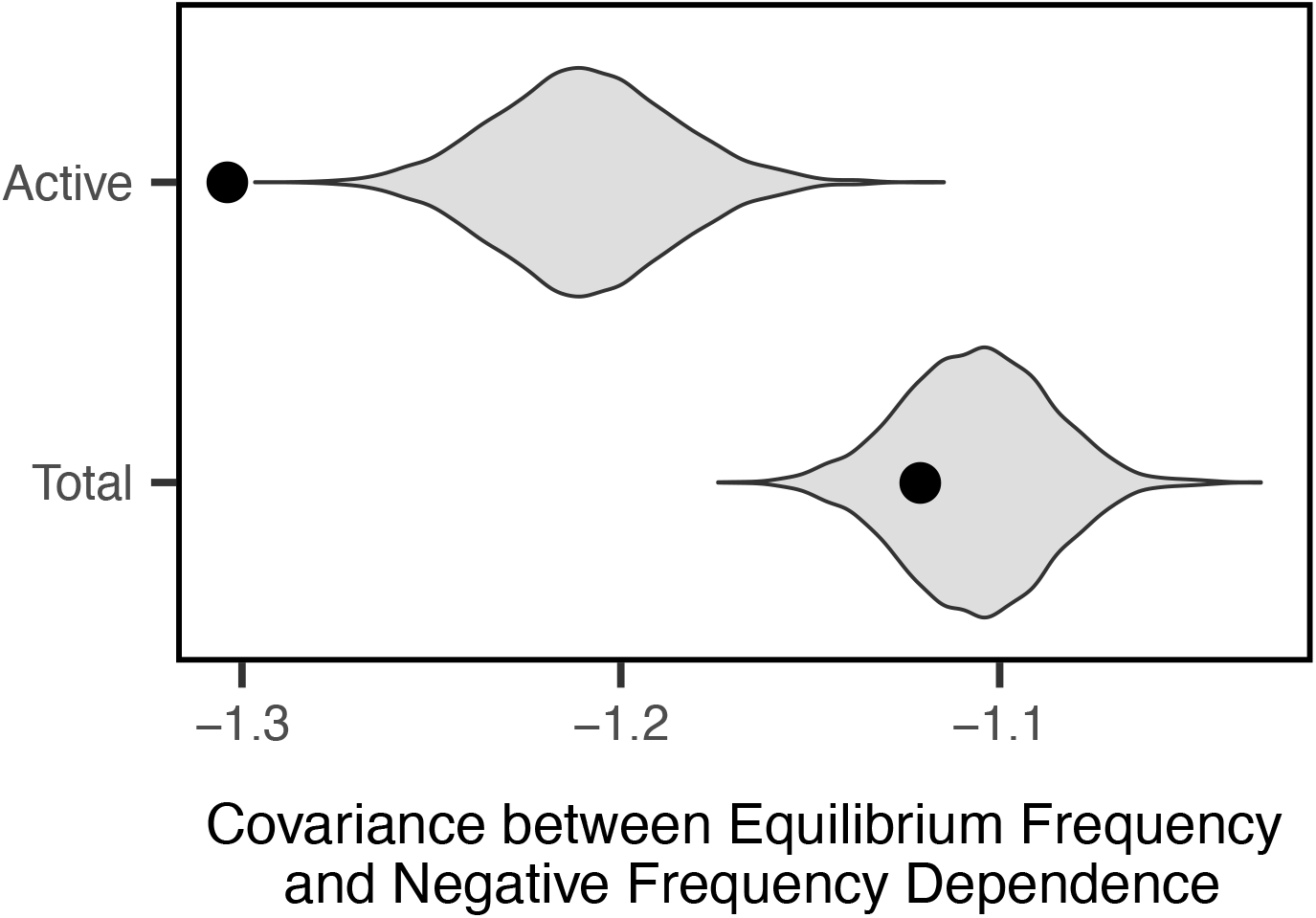
Negative frequency dependence (NFD) among persistent bacterial taxa (n = 82) was significantly stronger for rare than common taxa only in the active portion of the community. The degree of asymmetry in NFD is determined by the covariance between the equilibrium frequency of each OTU and its strength of NFD; negative covariance indicates that rarer taxa exhibit stronger NFD. Compared with expected covariances from a null distribution, the standardized effect size (SES) of the observed covariance in the active portion of the community was −4.03, while the SES of the total community was −0.77. The overall strength of NFD (observed NFD / mean NFD) was 1.08 in the active portion and 1.01 in the total community. The metabolic state on the y-axis indicates whether the NFD comparison is for the active portion of the community (inferred from 16S rRNA transcripts) or the total portion (inferred from 16S rRNA genes, i.e., DNA).

### Seed bank dynamics

Our data suggest seed banks of dormant or slow growing individuals contribute to the maintenance of diversity. Over the course of our study, total richness ranged from 1.2–2.0 times higher than the richness of the active portion of the community (Fig. 4). Furthermore, this discrepancy between total and active richness exhibited seasonality, demonstrating a time-varying role for the bacterial seed bank. In particular, the seed bank played a weaker role (i.e., active and total richness were more similar in magnitude) during the summer, while proportionally higher diversity was found in the seed bank over winter, when growing conditions may be less optimal (Fig. 4). In addition, the taxa that exhibited more reactivations from the seed bank were also the taxa that were, on average, consistently rare when active in the community (Fig. 5). In contrast, common taxa exhibited fewer transitions between active and inactive states in the community. Compared to null models of community dynamics, the observed relationship between relative abundance and reactivation actually implies a far stronger role for the seed bank among taxa of low-to-intermediate abundance ranks than expected by chance alone (Fig. S3), suggesting that the seed bank may be an important source of reestablishment in the active community. Thus, our findings support the view that rare taxa in the community benefit from life-history strategies such as slow growth or dormancy that minimize the probability of local extinction.

**Figure 4.**
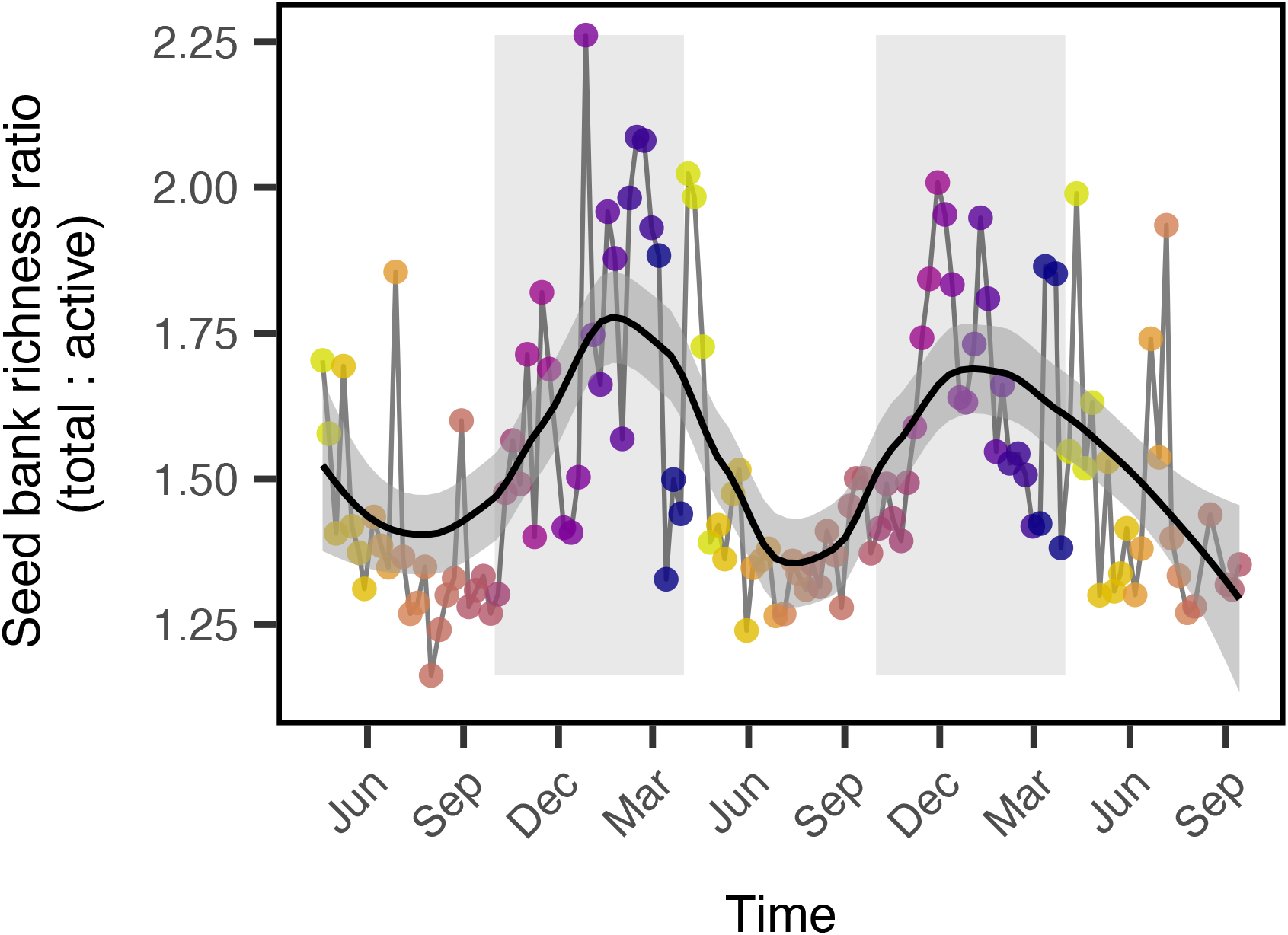
Seasonal importance of the seed bank for bacterial diversity in University Lake. Richness was much higher in the total community, relative to the active community, during the fall and winter months. The active and total communities converged over the summer, indicated by values on the y-axis closer to 1. Warmer colors correspond to spring and summer months, while cooler colors correspond to winter months. Shaded regions correspond to the fall and winter months (October through March).

**Figure 5.**
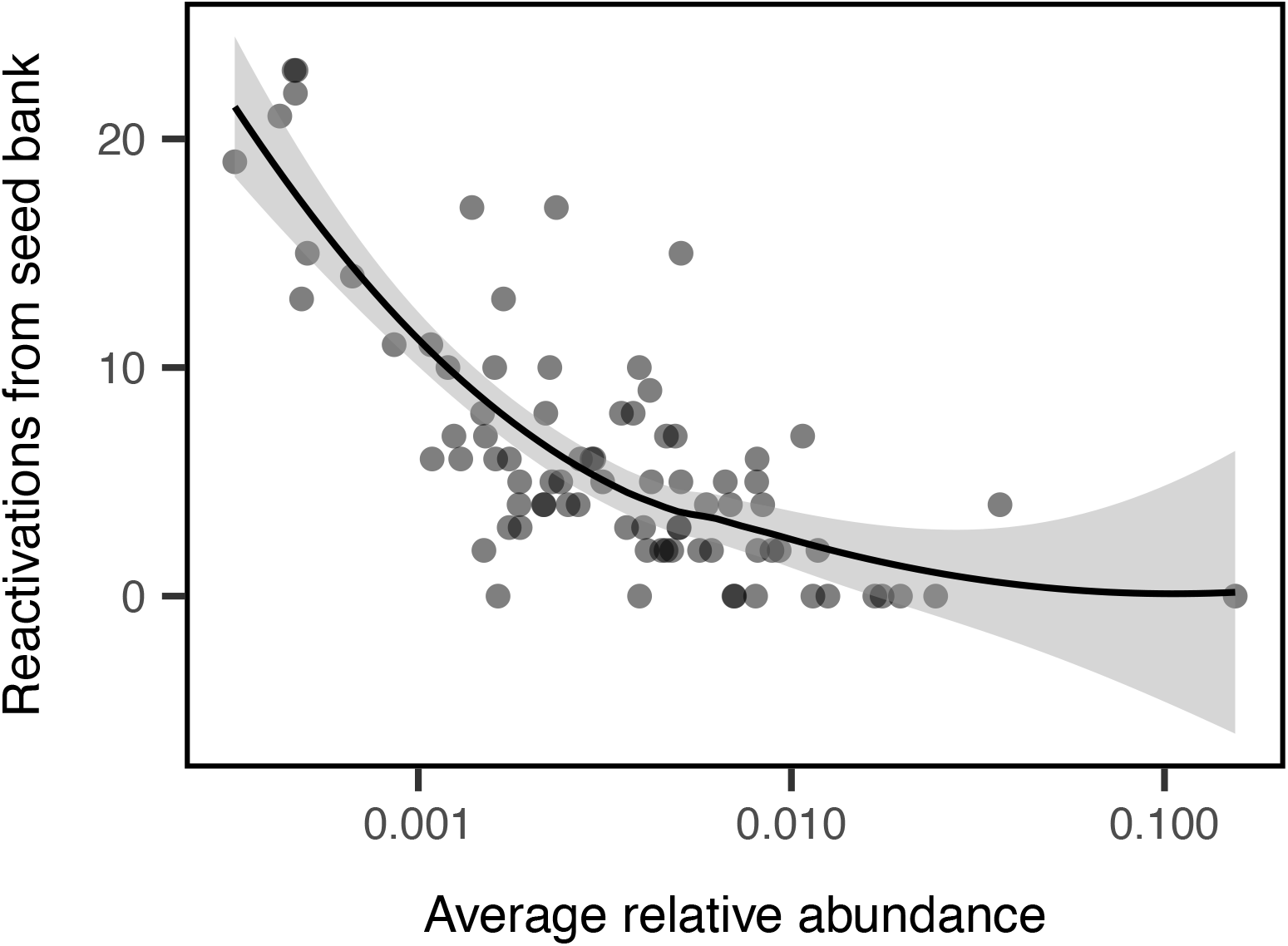
Rare taxa showed more seed bank transitions than common taxa. For the 82 persistent taxa identified over the time series, OTUs that were (on average) rare in the active portion of the community had a higher number of reactivations from the seed bank, while more common taxa had fewer reactivations. Regression lines are locally estimated scatterplot smoothing (LOESS).

## DISCUSSION

Our findings from a multi-year survey support the view that biodiversity was maintained by stabilizing mechanisms, including niche differentiation and seed bank dynamics, that generated negative frequency dependence (NFD) in a natural bacterioplankton community. High resolution sampling revealed recurrent seasonality in community dynamics (Fig. 1), driven by taxon-specific responses to annual environmental fluctuations (Fig. 2, Fig. S2). Our results also showed that the maintenance of diversity may be enhanced by life-history strategies, such as slow growth or dormancy, that buffered rare taxa from local extinction during environmentally unfavorable periods (e.g., winter) and facilitated reestablishment when conditions improved (Fig. 4–5). These apparent niche differences and seed bank dynamics contributed to stabilizing biotic interactions (e.g., stronger intra- than interspecific limitation) among rare, but metabolically active, taxa in the community (Fig. 3).

### Negative frequency dependence in microbial communities

We found evidence for stabilization through negative frequency dependence (NFD). While documented in some plant and animal assemblages (Harpole & Suding 2007; Yenni *et al.* 2017; Rovere & Fox 2019), observations of NFD in complex microbial communities are uncommon. In particular, our study revealed disproportionately strong NFD for rare taxa, offering an explanation for why some taxa appear to stably persist at low relative abundances in nature (Alonso-Sáez *et al.* 2015; Lindh *et al.* 2015), potentially as members of the “rare biosphere” (Sogin *et al.* 2006; Lynch & Neufeld 2015; Shade *et al.* 2018). Our approach also allowed us to identify stabilizing mechanisms operating among metabolically active rare taxa that were not detectable from the dynamics of the total bacterial community (Fig. 3). Ignoring this metabolic heterogeneity can obscure inferences of underlying ecological processes (Wisnoski *et al.* 2020), and would have gone otherwise undetected in this study as well.

Stronger NFD among rare taxa is also important for coexistence in plant and animal communities, but the magnitude of this effect varies across taxonomic groups (Yenni *et al.* 2017; Rovere & Fox 2019). For example, NFD is less asymmetric for herpetofauna than plant or mammal communities (Yenni *et al.* 2017), possibly due to higher evenness (Rovere & Fox 2019). Compared to macro-organismal systems, the degree of NFD asymmetry in our highly uneven bacterial community was moderate (SES = −4.03, covariance ratio = 1.08), suggesting that coexistence among rare taxa may be weak, or that additional factors not captured by this metric are important for maintaining diversity in our study system. However, we provide critical evidence that active bacteria mediate the biotic interactions responsible for generating stronger NFD among rare taxa in the community. Consistent with prior work showing that rare taxa may be disproportionately active in freshwater bacterial communities (Jones & Lennon 2010), our study demonstrates that rare, metabolically active bacteria may also be critical for the long-term maintenance of bacterial diversity.

### Dynamic microbial seed banks

Seed bank dynamics are thought to maintain diversity in fluctuating environments. In particular, seed banks provide a demographic buffering effect that satisfies one criterion of the storage effect. Evidence for coexistence via the storage effect comes largely from communities of desert annuals (Pake & Venable 1996; Angert *et al.* 2009), grasslands (Adler *et al.* 2006), tropical trees (Usinowicz *et al.* 2012), zooplankton (Cáceres 1997), and marine fish (Secor 2007). In most bacterial studies, the role of seed banks for coexistence has been inferred from short-term observations, but here we provide temporal evidence that bacterial seed banks may be important for community dynamics and the maintenance of diversity over longer, multi-annual time scales. In the temperate climate of our study lake, different taxa showed maximum growth rates at different times of the year, coinciding with seasonal transitions in environmental conditions (Fig. 2), and contrasting active and total community dynamics suggested buffered population dynamics (Figs. 3–5). Bacterial taxa may also exhibit more fine-grained differences in temporal niches than can be characterized by peak activity, which may be a somewhat conservative estimate of temporal niche differences. Nevertheless, our data provide evidence that two out of the three criteria for a storage effect (differential responses to the environment and buffered population dynamics) may be operating in the community.

The third criterion of the storage effect is that there is covariance between environmental conditions and competition. Documenting this pattern, especially in highly diverse communities, can often be a challenge. Our study demonstrated that species experienced greater self-limitation (consistent with stronger intraspecific than interspecific competition) when they were more common in the active community (and were thus more likely favored by the environment) (Fig. 3, Fig. S2), and diversity was maintained during potentially unfavorable growth environments (Fig. 4), but it is unclear whether environmental fluctuations in this system generate the covariance between environment and competition necessary for a storage effect (Chesson 2000; Miller & Klausmeier 2017). Theoretical models indicate that the storage effect may be more likely to evolve when species’ generation times are much shorter than the timescale of environmental fluctuations (Miller & Klausmeier 2017), a scenario that is well aligned with bacterioplankton living in a highly seasonal north temperate lake. In addition, we cannot rule out the potentially strong contribution to NFD by another non-mutually exclusive class of fluctuation-dependent mechanisms. Namely, relative nonlinearity in competition can promote coexistence if species differ in their responses to competition in ways that benefit their competitors (Yuan & Chesson 2015; Letten *et al.* 2018; Hallett *et al.* 2019). While our data cannot provide definitive proof, the documented patterns are consistent with the criteria needed for a storage effect to contribute to the long-term maintenance of bacterial diversity.

### Seasonal reoccurrence in bacterial communities

Seed banks may also have more general implications for bacterial community dynamics. The persistence of taxa with temporal niche differences could contribute to the repeatability of summer community dynamics in the active portion of the community (Hellweger *et al.* 2008) by favoring overwinter survival (Fig. 4). For example, we found that the seed bank exhibited seasonality, such that diversity stored in the seed bank was maximized when environmental conditions (e.g., water temperature, resource/consumer densities) were least favorable for bacterial growth (Neuenschwander *et al.* 2015). This pattern is consistent with the notion that dormant seed banks help buffer individuals from harsh conditions. In addition, transitions from inactive to active metabolic states were more frequently detected among taxa that were, on average, rare when active in the community.

Analogous to the methodological challenges of finding and identifying dormant individuals in non-microbial seed banks (e.g., plants), detection limits may affect the classification of metabolically active bacteria. Nevertheless, our reactivation metric should capture rapid shifts in the metabolically active portion of the community. Indeed, when compared to null models, our observations indicate that bacteria of rare-to-intermediate abundance ranks exhibited more frequent reactivations than would be expected by chance alone (Fig. S3), providing further evidence that seed banks are likely important sources of recolonization for bacterial communities inhabiting seasonal freshwater environments. Overall, these patterns suggest that recurrent environmental cues regulate active community dynamics by favoring different taxa at different times of the year, and that seed banks are important for maintaining these seasonal community trajectories at multi-annual timescales.

### Future directions and conclusions

Our study provides empirical evidence consistent with the theory that niche differences and seed bank dynamics stabilize bacterial communities and maintain diversity in nature. In the naturally fluctuating lake environment of our study, we demonstrated key differences in the diversity, dynamics, and stabilization between the active and total subsets of the bacterial community, but an ultimate goal is to tighten the mechanistic links between rates of ribosomal RNA transcription and *in situ* growth rates for individual taxa (Newton & Shade 2016; Papp *et al.* 2018) or through other techniques that involve the physical sorting of cells based on metabolic activity prior to sequencing (Couradeau *et al.* 2019; Reichart *et al.* 2020). While our results showed that stabilizing mechanisms generated NFD in the community, an important next step is to quantify the strengths and directions of the multiple fluctuation-independent and - dependent coexistence mechanisms that may be operating in diverse microbial communities (Letten *et al.* 2018; Ellner *et al.* 2019; Hallett *et al.* 2019).

A grand challenge at the intersection of microbial and community ecology is to extend the experimental investigations of microbial coexistence in the lab (Zhang *et al.* 2010; Letten *et al.* 2018) into systems reflecting the high diversity and complex interaction networks of most natural microbial communities. It will require careful experimentation as well as a clear consideration of the spatial scale of the study (e.g., to account for sampling biases and immigration that deviate from clear alignment with coexistence theory). For example, it may also be important to consider the dispersal of terrestrial bacteria into aquatic ecosystems (Crump *et al.* 2012), since immigration could contribute to inferences made about local processes. However, previous work in our study system revealed that most terrestrial bacteria were metabolically inactive, most likely reflecting the abrupt environmental transitions that accompany cross-ecosystem dispersal (Wisnoski *et al.* 2020). Thus, the lack of asymmetric NFD we observed in the total community could also arise in part from allochthonous inputs of inactive bacteria that decouple local population growth of each OTU from its relative abundance in the community. However, by focusing on the dynamics of metabolically active bacteria, our approach was able to uncover the presence of stabilizing biotic interactions that may have otherwise been obscured by metabolic heterogeneity in the total community.

In conclusion, we show that stabilizing biotic interactions and the ability to engage in dormancy or slow growth strategies play important roles in maintaining microbial diversity in a natural ecosystem over a multi-year time scale. Our results demonstrate the mechanisms at the community scale that preserve Earth’s vast microbial diversity, building on other explanations that emphasize the importance of metabolic diversity (Sala *et al.* 2008), capacity for rapid growth (Shade *et al.* 2014), and spatial scale (Vos *et al.* 2013). In particular, strong NFD offers a new explanation for why the majority of bacterial taxa persist at low average relative abundances in nature (Lynch & Neufeld 2015). Furthermore, our work builds on inferences about the roles of microbial dormancy (and other persistence strategies) obtained from shorter time scales, and provides temporal evidence that dormancy is an important buffer against local extinction over longer time scales. More generally, our work demonstrates the importance of stabilization in microbial systems, offering new insight into the long-term maintenance of microbial diversity.

## ACKNOWLEDGEMENTS

We acknowledge technical support from Brent Lehmkuhl and Drew Meyer. This work was supported by the National Science Foundation (DEB-1442246 and 1934554 to JTL), US Army Research Office Grant (W911NF-14-1-0411 to JTL), National Aeronautics and Space Administration (80NSSC20K0618 to JTL), and the Department of Biology at Indiana University (Louise Constable Hoover Fellowship to NIW). Data and code for the project can be found at NCBI (BioProject PRJNA664410) and a Zenodo archive of the GitHub repository (https://github.com/LennonLab/ul-seedbank).

## SUPPLEMENTAL INFORMATION

**Table S1.** Operational taxonomic units (OTUs) that were classified as persistent in the bacterioplankton community based on being detected in ≥80% of the total (i.e., DNA) community samples. The table is sorted by Julian date of max growth.

| OTU | Class | Max growth rate ($d^{-1}$) | Date of max growth |
| --- | --- | --- | --- |
| Otu00034 | Alphaproteobacteria | 0.3426 | 2014-01-17 |
| Otu00045 | Betaproteobacteria | 0.8527 | 2014-01-03 |
| Otu00196 | Actinobacteria | 0.5873 | 2015-01-09 |
| Otu00019 | Cytophagia | 0.7346 | 2014-02-14 |
| Otu00039 | Betaproteobacteria | 0.7167 | 2014-02-14 |
| Otu00102 | Betaproteobacteria | 0.6729 | 2015-02-28 |
| Otu00105 | Alphaproteobacteria | 0.8666 | 2014-02-28 |
| Otu00065 | Sphingobacteriia | 0.8563 | 2014-03-21 |
| Otu00292 | Alphaproteobacteria | 0.6851 | 2014-03-07 |
| Otu00006 | Sphingobacteriia | 0.3643 | 2013-04-25 |
| Otu00012 | Betaproteobacteria | 0.8784 | 2014-04-18 |
| Otu00014 | Actinobacteria | 0.4906 | 2015-04-26 |
| Otu00016 | Actinobacteria | 0.7836 | 2014-04-18 |
| Otu00017 | Actinobacteria | 1.0200 | 2015-04-04 |
| Otu00021 | Gammaproteobacteria | 1.0449 | 2014-04-18 |
| Otu00033 | Alphaproteobacteria | 0.5306 | 2014-04-25 |
| Otu00048 | Verrucomicrobiae | 0.7426 | 2015-04-11 |
| Otu00049 | Actinobacteria | 0.6090 | 2014-04-04 |
| Otu00055 | Flavobacteriia | 0.7950 | 2014-04-18 |
| Otu00058 | Armatimonadia | 0.8153 | 2015-04-11 |
| Otu00148 | Bacteria sp. | 0.7646 | 2013-04-25 |
| Otu00172 | Gammaproteobacteria | 0.6729 | 2015-04-11 |
| Otu00219 | Betaproteobacteria | 0.7503 | 2014-04-18 |
| Otu00002 | Actinobacteria | 0.3190 | 2015-05-03 |
| Otu00008 | Actinobacteria | 0.3637 | 2013-05-09 |
| Otu00022 | Opitutae | 0.8003 | 2015-05-03 |
| Otu00031 | Cytophagia | 0.6593 | 2014-05-09 |
| Otu00051 | Flavobacteriia | 0.9187 | 2013-05-09 |
| Otu00062 | Flavobacteriia | 0.7836 | 2015-05-23 |
| Otu00064 | Alphaproteobacteria | 0.6964 | 2013-05-29 |
| Otu00113 | Bacteroidetes sp. | 0.7950 | 2013-05-09 |
| Otu00116 | Betaproteobacteria | 0.8373 | 2014-05-09 |
| Otu00151 | Betaproteobacteria | 0.7259 | 2013-05-17 |
| Otu00183 | Bacteria sp. | 0.6090 | 2015-05-03 |
| Otu00200 | Bacteria sp. | 0.6851 | 2015-05-23 |
| Otu00083 | Flavobacteriia | 1.0031 | 2015-06-06 |
| Otu00087 | Betaproteobacteria | 0.5873 | 2014-06-05 |
| Otu00098 | Betaproteobacteria | 0.8104 | 2013-06-14 |
| Otu00123 | Sphingobacteriia | 0.7259 | 2014-06-20 |
| Otu00194 | Deltaproteobacteria | 0.9277 | 2014-06-13 |
| Otu00294 | Alphaproteobacteria | 0.6964 | 2013-06-21 |
| Otu00004 | Actinobacteria | 0.3500 | 2015-07-11 |
| Otu00009 | Gammaproteobacteria | 1.3329 | 2013-07-26 |
| Otu00011 | Betaproteobacteria | 0.9841 | 2015-07-18 |
| Otu00192 | Bacteria sp. | 0.4350 | 2013-07-26 |
| Otu00195 | Actinobacteria | 0.6277 | 2014-07-18 |
| Otu00020 | Betaproteobacteria | 0.4150 | 2013-08-01 |
| Otu00026 | Betaproteobacteria | 0.6277 | 2013-08-01 |
| Otu00029 | Actinobacteria | 0.6593 | 2013-08-23 |
| Otu00036 | Alphaproteobacteria | 0.7576 | 2013-08-16 |
| Otu00037 | Actinobacteria | 0.7646 | 2014-08-29 |
| Otu00052 | Alphaproteobacteria | 0.6444 | 2013-08-09 |
| Otu00076 | Actinobacteria | 0.6444 | 2014-08-08 |
| Otu00112 | Alphaproteobacteria | 0.5873 | 2014-08-23 |
| Otu00226 | Opitutae | 0.6729 | 2015-08-02 |
| Otu00250 | Actinobacteria | 0.5306 | 2013-08-23 |
| Otu00010 | Proteobacteria sp. | 0.6277 | 2014-09-26 |
| Otu00024 | Bacteroidetes sp. | 0.7259 | 2014-09-19 |
| Otu00056 | Cytophagia | 0.7711 | 2013-09-06 |
| Otu00067 | Betaproteobacteria | 0.5306 | 2015-09-02 |
| Otu00177 | Proteobacteria sp. | 0.5306 | 2013-09-20 |
| Otu00005 | Sphingobacteriia | 0.5294 | 2014-10-04 |
| Otu00015 | Actinobacteria | 0.6277 | 2014-10-17 |
| Otu00018 | Gammaproteobacteria | 0.9516 | 2014-10-11 |
| Otu00060 | Betaproteobacteria | 1.0031 | 2013-10-25 |
| Otu00066 | Betaproteobacteria | 0.8153 | 2013-10-25 |
| Otu00082 | Bacteroidetes sp. | 0.8003 | 2014-10-04 |
| Otu00154 | Alphaproteobacteria | 0.7070 | 2014-10-04 |
| Otu00158 | Gammaproteobacteria | 0.9141 | 2013-10-04 |
| Otu00001 | Betaproteobacteria | 0.2860 | 2013-11-15 |
| Otu00007 | Betaproteobacteria | 0.4577 | 2013-11-15 |
| Otu00038 | Actinobacteria | 0.7346 | 2014-11-29 |
| Otu00047 | Betaproteobacteria | 0.6729 | 2013-11-15 |
| Otu00073 | Betaproteobacteria | 0.4906 | 2013-11-22 |
| Otu00077 | Flavobacteriia | 0.7774 | 2013-11-15 |
| Otu00109 | Actinobacteria | 0.6729 | 2013-11-15 |
| Otu00118 | Actinobacteria | 0.6090 | 2013-11-22 |
| Otu00129 | Alphaproteobacteria | 0.4906 | 2013-11-15 |
| Otu00198 | Betaproteobacteria | 0.9210 | 2013-11-15 |
| Otu00208 | Betaproteobacteria | 0.6964 | 2013-11-22 |
| Otu00217 | Proteobacteria sp. | 0.7167 | 2013-11-22 |

**Figure S1.**
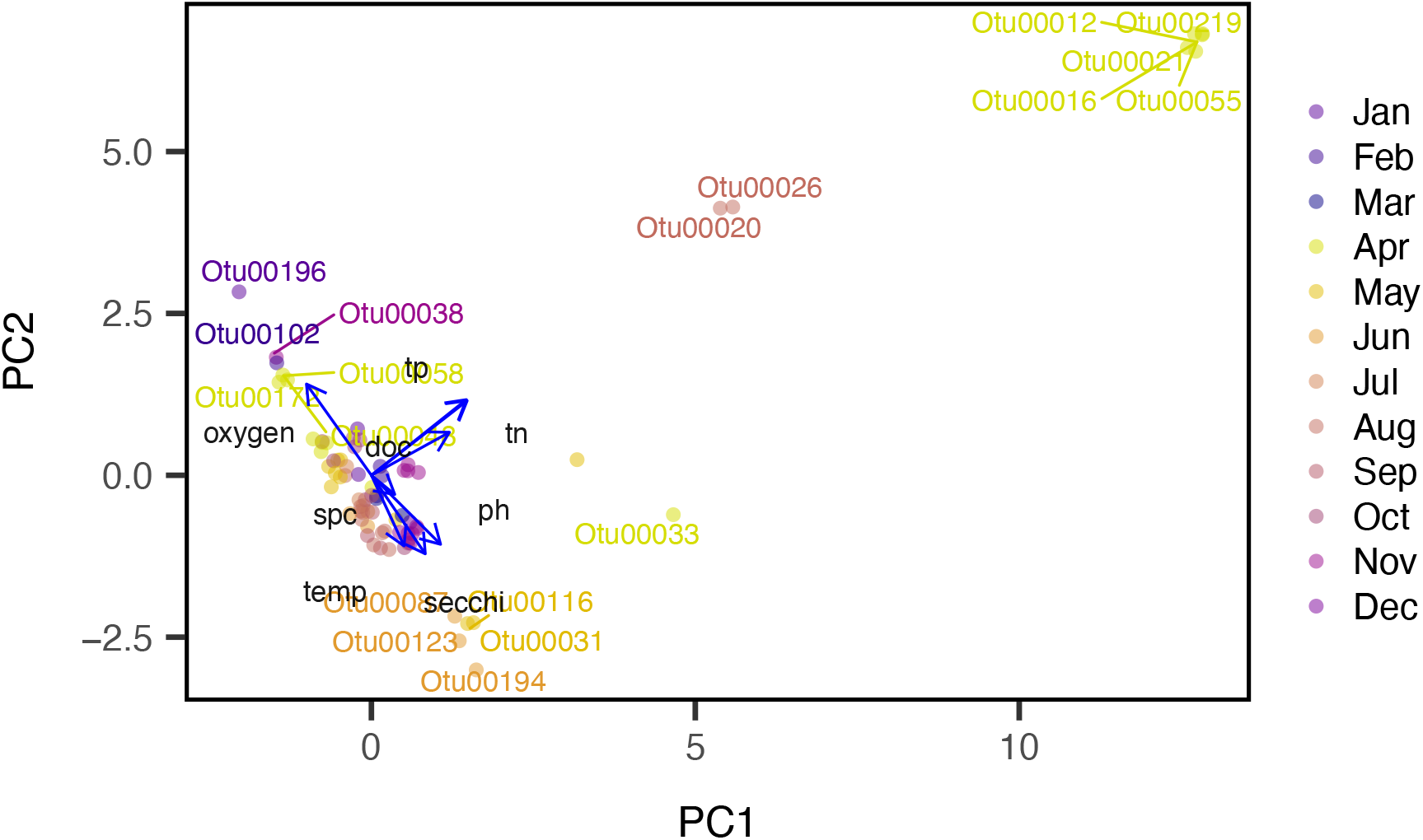
Temporal partitioning of maximum growth rates corresponds to environmental conditions. Here, we depict a principal component analysis of the environmental conditions at each date. OTU labels indicate the individual taxa that exhibit maximum growth at the corresponding time of the year. Vectors indicate the loadings of environmental variables along the two PC axes. Warmer colors correspond to spring and summer months, cooler colors correspond to winter months.

**Figure S2.**
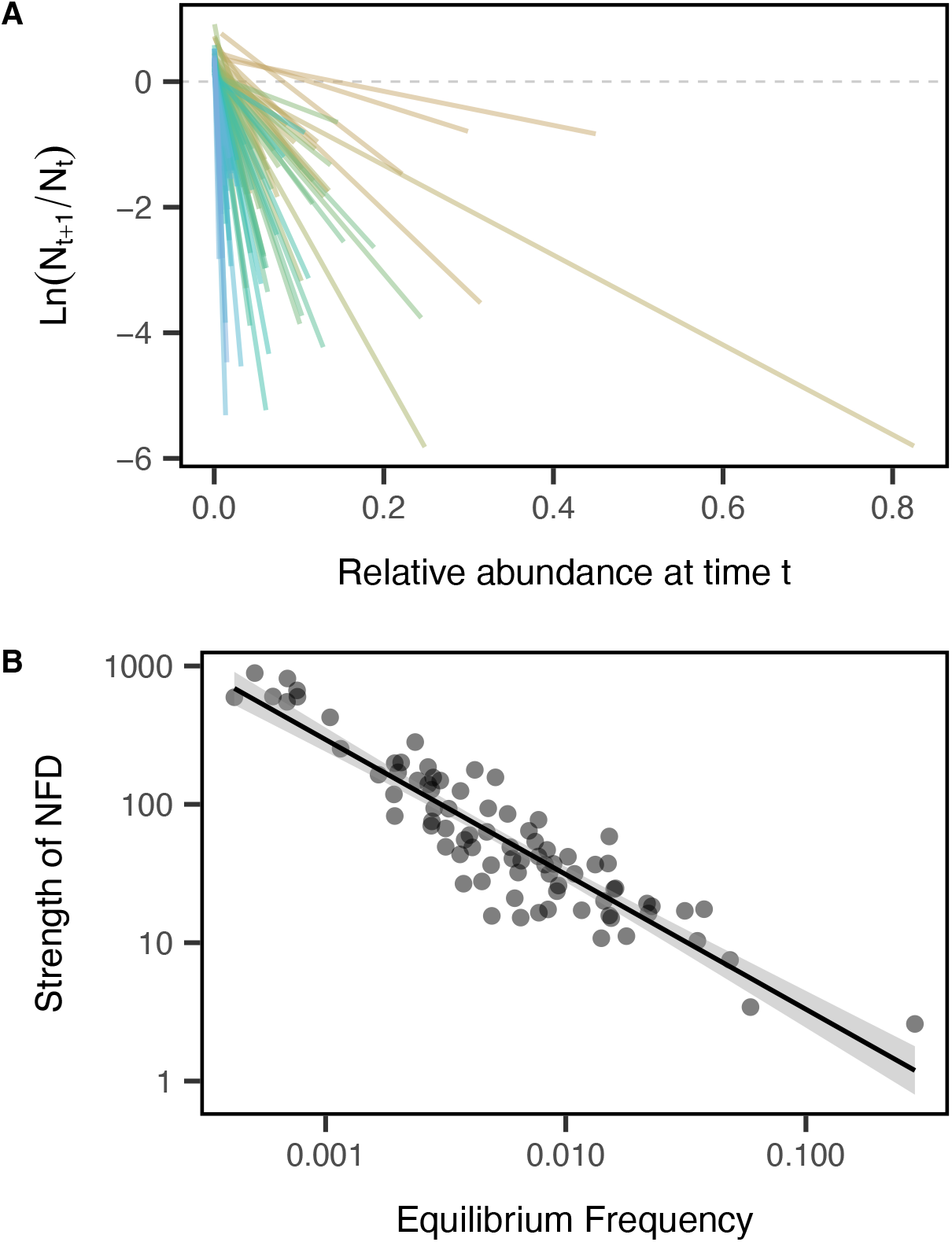
Negative frequency dependence (NFD) in the active portion of the community for the 82 persistent bacterial taxa. (A) Relationship between the rate of change of an OTU and its relative abundance. Depicted in this graph are simple linear regression fits for the 82 taxa individually, data points not shown to reduce clutter. Negative relationships indicate NFD growth and variation in slopes indicates variation in the strength of NFD. (B) Rare taxa (lower equilibrium frequencies) exhibit stronger NFD, while common taxa (higher equilibrium frequency) have weaker NFD.

**Figure S3.**
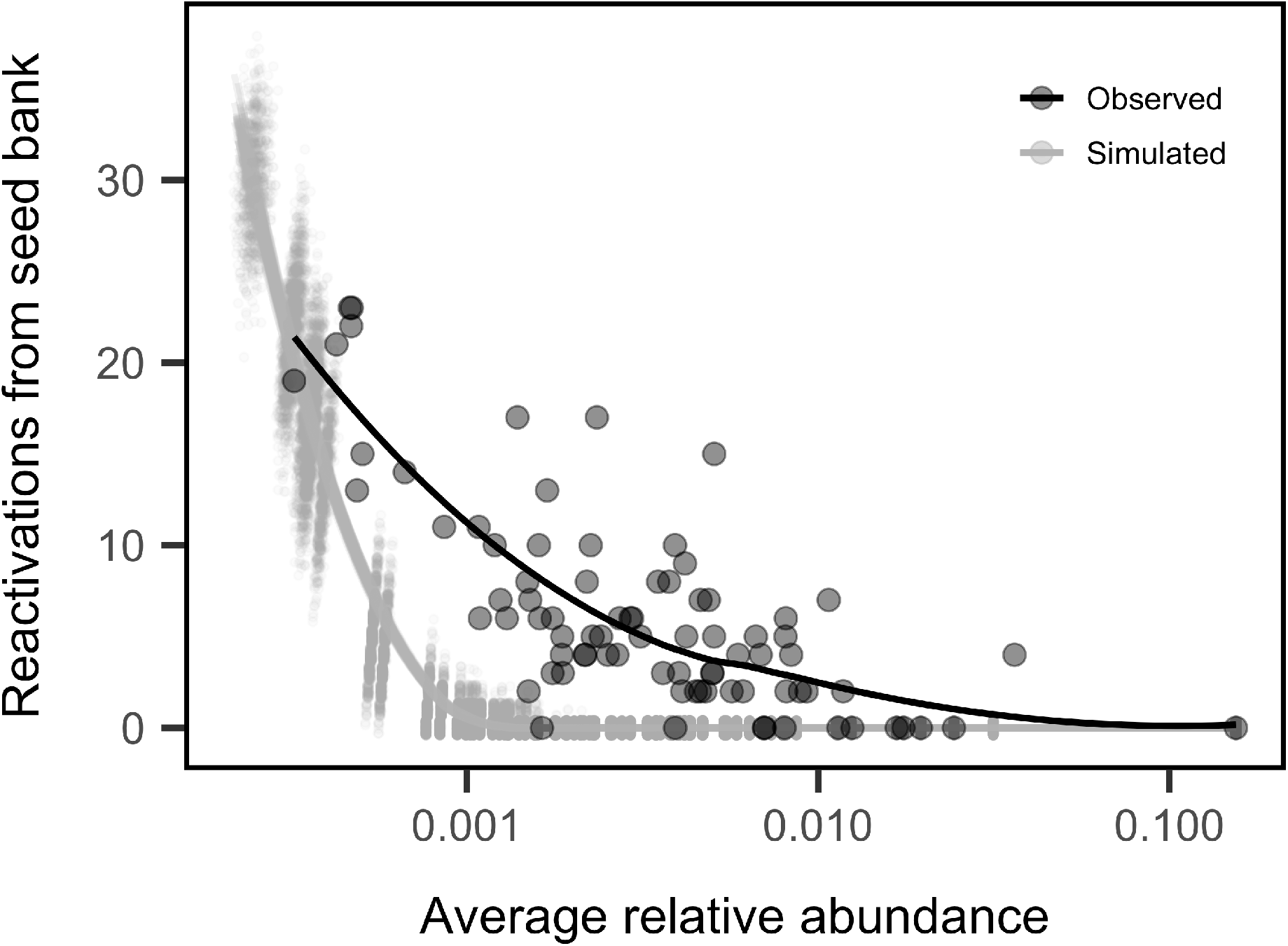
Observed reactivations from the seed bank are more frequent than expected by chance alone. Null model simulations (n = 1000) describe the range of expectations expected by chance for the relationship between relative abundance and reactivation rate (gray points). For observed data (black points) and the null simulations, we fit local regression lines with locally estimated scatterplot smoothing (LOESS).

